# Algebraic structures emerge from the self-supervised learning of natural sounds

**DOI:** 10.1101/2024.03.13.584776

**Authors:** Pierre Orhan, Yves Boubenec, Jean-Remi King

**Affiliations:** Laboratoire des systèmes perceptifs, Département d’études cognitives, École normale supérieure, PSL University, CNRS, 75005 Paris, France; Meta, Paris, France

**Keywords:** Self-supervised learning, Statistical learning, Language, Music, Structure

## Abstract

Humans can spontaneously detect complex algebraic structures. Historically, two opposing views explain this ability, at the root of language and music acquisition. Some argue for the existence of an innate and specific mechanism, like “merge” (Chomsky) or “neural recursion” (Dehaene). Others argue that this ability emerges from experience (e.g. Bates): i.e. when generic learning principles continuously process sensory inputs. These two views, however, remain difficult to test experimentally. Here, we use deep learning models to evaluate the factors that lead to the spontaneous detection of algebraic structures in the auditory modality. Specifically, we train multiple deep learning models with a variable amount of natural sounds and a self-supervised learning objective. We then expose these models to the experimental paradigms classically used to evaluate the processing of algebraic structures. Like humans, these models spontaneously detect repeated sequences, probabilistic chunks and complex algebraic structures. Also like humans, this ability diminishes with structure complexity. Importantly, this ability can emerge from experience alone: the more the models are exposed to natural sounds, the more they spontaneously detect increasingly complex structures. Finally, this ability does not emerge in models pretrained only on speech, and emerges more rapidly in models pretrained with music than environmental sounds. Overall, our study provides an operational framework to clarify sufficient built-in and acquired principles that model human’s advanced capacity to detect algebraic structures in sounds.

**Significance Statement:** Experimentalists have repeatedly observed a human advantage in the detection of algebraic structures, notably through auditory paradigms. This ability to detect structure is thought to be key to the emergence of complex cognitive operations. Yet, it remains debated if this ability is discovered or innate in the form of a specific mechanism. In this article, the authors show how a model progressively and spontaneously learns to detect auditory structure. The model replicate several experimental findings but only under certain developmental conditions. Notably, exposition to music or environmental sounds, but not speech, is sufficient for the emergence of algebraic structure detection. As a result, this work proposes self-supervised learning as a developmental model of abstract cognitive abilities.

## Introduction

### The controversial underpinning of structure building in the human mind

Humans have a unique ability to detect symbolic structures: Unlike other mammals, we spontaneously detect syntax from speech input (1–5), and geometrical symmetries from simple drawings (6, 7). To account for this singular ability, linguistics, artificial intelligence and neuroscience communities have explored two competing views: “Rationalists” argue that an innate and specific mechanism, like “Merge” (8) or “a recursive neural code” (7), allows the human brain to instinctively represent information as tree structures. On the other hand, “Empiricists” argue that this cognitive ability need not require a specific, nor an innate mechanism, but rather emerges from an efficient statistical learning of naturalistic inputs (9).

### Brain and behavioral bases of minimal structure building

To confront these perspectives, a variety of minimalist protocols focusing on the brain and behavior’s fundamental aspects of structure building have been proposed (10–15). These protocols typically use artificial stimuli to ensure that participants are not already familiar with the tested structures. In the auditory domain in particular, Saffran et al. (10) observed that children as young as eight-month-old spontaneously segment recurring sequences of syllables. In adults, Al Roumi et al. demonstrated that the ability to detect auditory structures strongly depends on their algebraic complexity (16). These spontaneous behavioral abilities do not appear to be associated with a unique neural system: First, the violation of simple auditory regularities elicit an early mismatch negativity (MMN) (17) in electro-encephalography (EEG) recordings, while more complex auditory structures are associated with a late positivity (P300b) (18, 19). Second, the detection of long auditory sequences depends on a sustained, distributed and elevated Magneto-Encephalography (MEG) response (20). Finally, the complexity of auditory structures is linked to parietal activations, and seems dissociated from the language network in the brain (16).

### Modeling structure building

Consequently, a variety of neural or symbolic models have been proposed to account for these behavioral and neural observations. Local plasticity in the thalamo-cortical circuit has been proposed to account for MMN in auditory irregularities (21). Prediction by Partial Matching (PPM) models, like Information Dynamics Of Music (IDyOM) (22) model or PPM-decay model (23), have been shown to provide optimal predictions for melodic expectation, and to mirror human behavior in several tasks (24). In addition, a variety of neuro-symbolic models have been proposed to tackle systematic compositions of input sequences (25–29). The common – although not always explicit – denominators across these efforts, is the idea that discovering the specific mechanism responsible for spontaneously detecting structures would prove critical to build system that act and thinks like humans.

### Main challenge

Overall, however, these efforts face a common challenge: all models are crafted to operate exclusively on one experimental protocol. This approach thus presents two major limitations. First, no model has been systematically tested on its ability to account for a large variety of experimental observations. Second, no model actually leverage the possibility to learn structures from natural stimuli. Detecting artificial structures with such a restricted approach thus constrains us to either design a hard-wired system, or to extensively train it with our artificial stimuli. In sum, the rationalist view benefits from an intrinsic head-start.

### Modern AI to the rescue of empiricists

Here, we argue for an alternative approach. If structure building emerges from learning, then we should investigate models that can learn from naturalistic stimuli, and then only test their ability to detect artificial structures – not the other way around. Following the recent observation that large language models (LLMs) demonstrate remarkable generalization abilities without additional training (30–32), we hypothesize that the representations modern AI systems can learn from naturalistic data may suffice to lead them to spontaneously build adequate structures when presented to artificial structures like algebraic sequences. To avoid LLMs, and their unreasonably large and symbolic datasets, we here focus on self supervised learning (SSL) architectures designed to process natural sounds. Previous works, in vision (33, 34) and audition (35), have shown that such architectures develop human-like representations from a similar data regime.

### Investigation

Specifically, we use a standard SSL architecture, Wav2vec2 (36), to train a series of deep neural networks to “unmask” natural sounds. We then present these models to four psycho-acoustic protocols developed for structure detection, namely: (10)’s syllable chunking paradigm, (20)’s RangReg paradigm, (18)’s Local-Global and (16)’s Algebraic Patterns paradigm. To compare these models to humans, we evaluate the models’ ability to spontaneously detect auditory structures by measuring their surprisal in response to each sound, and in particular when the auditory regularities violate the artificial structure. Critically, we systematically investigate how the type and the amount of pre-training impact on the models’ ability to spontaneously detect auditory structures. Overall, our results reveal the conditions in which an efficient SSL architecture exposed to natural sounds may suffice to account for humans’ spontaneous ability to detect algebraic sequences.

## Approach

We aim to evaluate whether a model trained with a reasonable amount of naturalistic sounds stimuli, learns latent representations that allows it to spontaneously detect the structures of variably-complex algebraic or probabilistic sequences.

We focus on the auditory modality, where the detection of algebraic structures has been more systematically investigated than the visual modality.

### Model

We study a self-supervised neural network: Wav2vec2 (36). The model takes as input sound waveform. The model is structured hierarchically with local filtering (convolutions) followed by global contextual processing (transformer) (Fig. 1B). The model objective is to predict masked part of the sounds. More precisely, acoustic latent computed by the convolutions layer are first randomly masked. Based on the surrounding context, the model then tries to change this mask latent vector back to the original vector ^1^. The model loss contrasts this reconstruction with the target and a set of other latents used as negatives.

**Fig. 1.**
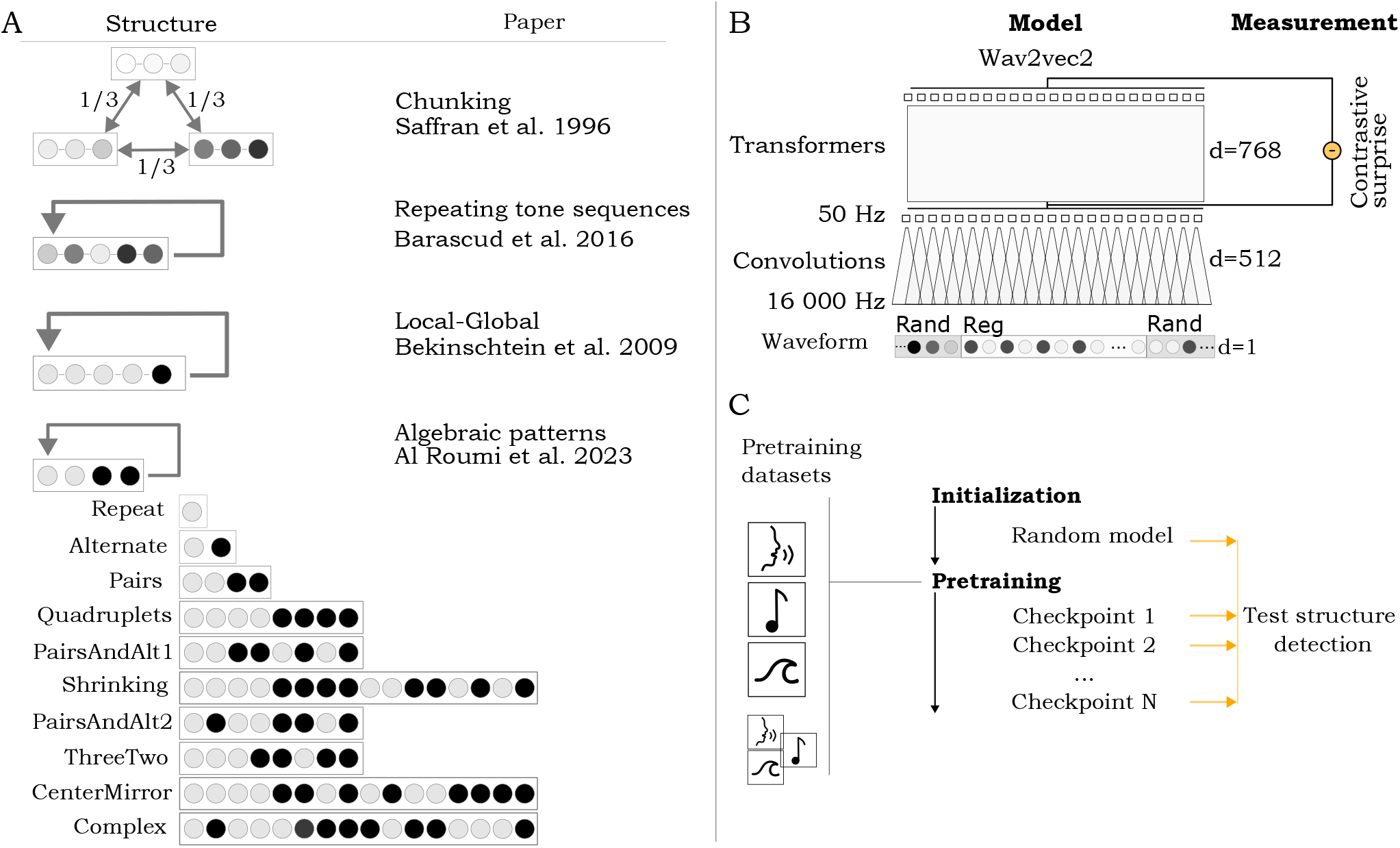
A: schematic of the structure of the sounds tested over the experiments. The right column indicate the experimental paper testing how humans detect the corresponding structure. B: schematic presentation of the Wav2vec2 model used. d refers to the dimensionality of the features at different layer. The model takes waveform as input so d=1 at the beginning. We measure the model’s surprise as the contrastive loss to each sound elements in the sound. This loss is measured by masking 20 ms (50Hz) latent vectors whose receptive field overlap with the sound element. This operation is repeated to get the model surprise of each sound element. C: Models are initially pretrained on one dataset among four possible: one for speech - Librispeech (37), one for music - FMA (38), one for environmental sounds - Audioset (39) deprived of speech and music and one that combine a subset of those. To see if and when structure detection emerge, we test the models on many checkpoints throughout the pretraining.

### Pretraining

The model is exposed (pretraining) before the experiments to several epochs of a modest (900 hours) sound dataset and optimised to diminish its contrastive loss^2^ by predicting masked latent acoustics for a maximal amount of 100,000 gradient steps. We first pretrain the model on a general audio dataset, composed of 3 × 300 hours of public datasets: Librispeech (speech, (37)), FMA (music, (38)) and Audioset (environmental sounds (39), we removed musical and speech sounds from this dataset thanks to the accompanying taxonomy). To investigate the effect of each dataset, we then pretrain the model separately on the full individual public datasets (about 900h each). The model is optimized with the original, base, parameters.

### Experimental paradigms

#### Protocols

We explore four experimental protocols (Fig. 1 A)(10, 16, 18, 20) investigating the behavioral and brain foundations of structure detection. These experiments assess subjects’ perception of structured sound sequences, exposing them to regular, random or deviant sequences. For simplicity, we standardize this variety of experimental protocols as follows. First, the pretrained model is exposed to random sounds. This allows us to evaluate its ‘basal’ surprise values. Second, we present it to a repeated structure (i.e. a regular sequence, generated from a subset of a larger alphabet), which allows us to verify the diminution of surprise. Finally, we present a random sequence to verify that the change in surprise is not just due to time (e.g. fatigue). The sound of this ending random sequences are drawn from the same set as the regular sequence, controlling for the effect of a diminishing alphabet, and instantaneously violating the preceding regular structure. We generate each rand-regular-rand sequences as a single waveform and present the whole sequence to our models.

#### Evaluation

To reveal if the model detects the regularity, we compute the model surprise on each sound element (tones or syllables) (Fig. 1B). The surprise is measured as the contrastive loss of the model prediction in response to the masking of each sound elements. Note that the model is consequently evaluated on the same audio file as many times as there are sound elements in the file, with the mask positioned each time over a different, probed, sound elements.

The model is evaluated in a “zero-shot” scenario, where it is presented with sound sequences but can’t optimize anymore its parameters.

## Results

### Statistical chunking of words

We tested if pretrained models could spontaneously discover words in a syllable stream. We replicated in-silico the experiment of Saffran et al. (10), which demonstrated that 8-month-old children can rapidly (less than 2 minutes) learn to detect 3-syllable words in a stream of syllables. A random stream of syllables was followed by a regular stream of syllables composed of 4 randomly alternating 3-syllable words (Fig. 2 A).

**Fig. 2.**
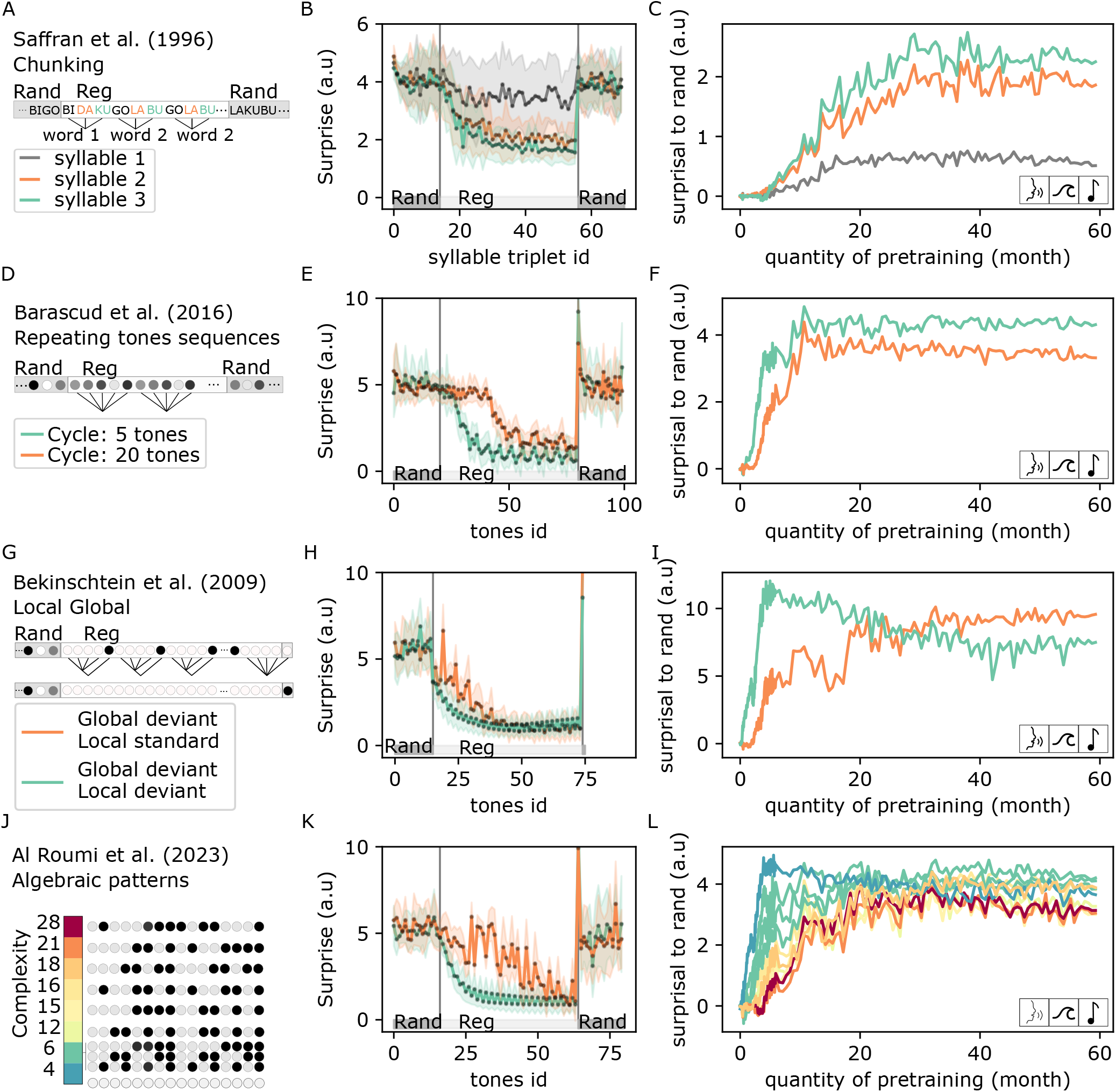
A-C) Zero-shot emergence of word chunking. **A**. Example of a syllable stream tested. **B**. Model contrastive loss to the first, second and third syllable is measured for each triplet of syllables (words) in a syllable stream. The loss is averaged over 30 trial of the tasks. The loss standard deviation across these trial is indicated as shaded area. (see supp. Fig. 4 for zooms). **C**. Evolution of the model ability to detect regular words as a function of its pretraining. This ability is measured by the difference between the mean contrastive loss of the second random sequence and the last repetition of the regular sequence. **D-E-F**. Same as A-B-C but with repeated tones sequences of cycle size 5 or 20 tones. **G-H-I**. Same as A-B-C but with a nested algebraic pattern from the Local Global paradigm. **J-K-L**. Same as A,B,C but with a set of 10 algebraic patterns of increasing complexity. In panel K, the “alternate” and “center-mirror” sequences are plotted, of complexity respective 6 and 21.

### Structure detection

We measured the model surprise to each syllables of the stream without changing the model weights (i.e. frozen model). As expected, the model surprise was high on the first part of the stimuli, composed of randomly alternating syllables (Fig. 2 B). Critically, the models’ surprise dropped when the stream became regular and groups of three syllables repeated to form four possible “words”. Notably, the model loss dropped on the second and third syllables of each word (Fig. 2 B). The model loss remained high for the first syllables, indeed the first syllables should not be predictable as words are randomly alternating in the syllable stream.

### Learning dynamic

We then investigated the learning dynamic of this ability. We repeated the experiments for 371 checkpoints spanning the models pretraining. We measured the difference between the model surprise during the regular stream, containing syllables grouped by words, and the ending stream, containing random sequences of syllables (Fig. 2 C). Overall, the model surprise to the random stream increased quasi monotonously with the amount of pretraining. Together, these results demonstrate that a passive exposure to natural sounds led the model to spontaneously detect repeated chunks of syllables without any additional training or specific built-in mechanism.

### Controls

To investigate further the experiments of Saffran et al. (10), we also tested the models’ reaction to a deviant group of three syllables. The deviant was either a never seen non-word (i.e a sequence of syllables that had never been seen) or a word that had been seen (the last syllable of a word followed by the first two syllable of another word). Similar to the eight-month-old children tested in Saffran et al. (10), the model’s surprise peaked significantly to these two types of deviants (supplementary Fig 2). This confirms the model ability to perform in-context discovery of words in this particular protocol, as humans.

### Detection of repeated tones sequences

Since the model can detect repeated 3-syllable motifs, we questioned if this ability was restricted to artificial syllable streams or could extend to longer and more complex sequences. We leveraged a paradigm based on rapid sequences of tones developed by Chait and colleagues. Using this paradigm, Barascud et al. (20) showed that humans were able to optimally detect transitions from random to repeated sequences of 5, 10 or 15 tones and sub-optimally for 20 tones. To compare our model to humans in this ‘Repeating tones’ task, we adapted the syllable approach described above as follows. As in Barascud et al. (20), tones were randomly sampled from a set of 20 tones covering a logarithmic frequency range from 222 to 2,000 Hz. A random stream of tones was followed by a regular stream, which repeated a sequence of tones, and ended by a new random stream, alternating tones composing the repeated sequences (Fig. 2 D). We tested repeated sequences of length 5 and 20.

### Structure detection

The model was able to detect the transition from regular to random for all sequences irrespective of their length (Fig. 2 E): i.e. its surprise was greater on the first random deviant tones than on preceding tones.

### Learning dynamic

This regularity detection emerged rapidly during pretraining (Fig. 2 F): it developed over the first “year” of sound exposure, with the detection of short sequences detection preceding that of long sequences.

### Controls

To detect repeated tones, the model could use the difference of acoustic between the repeating and the surrounding random sequences. Two arguments can be made against this hypothesis. First, detection happened on the first random tone and therefore is not using the long-term integration necessary to detect changes in acoustics. Second, we found that the model detected a single-tone deviation in the repeating sequence (sensitivity to deviant, D’, of 2.5, see methods for computation of D’), indicating that the model is accurately tracking the exact sequence of tones, and not long-term acoustics.

As discussed in the original experimental paper (20), the transition from a random to a regular sequence can only be detected by a causal system after the second motif appearance. However, Wav2vec2 uses a bidirectional transformer and is therefore non-causal. The original Wav2vec2 (36) therefore detects the transition from RAND to REG at the first tone of the REG sequence (supp. Fig 2), which is, by definition, different from human. For all experiments in the paper, we made the model partially causal by preventing it to use future information (see methods). This change recovered a human-like regularity detection with the surprise diminishing during the second repetition only.

Based on this observation, we can state that the model is able to detect syllabic regularities, but also repetition of arbitrary tone sequences.

### Detection of global deviant in tone sequences

We then replicate the Local-Global paradigm which has been extensively studied both in humans (40–44) and animals (45, 46). In this paradigm, two 5-tone sequences are used: a XXXXX sequence (local standard) and a XXXXY sequence (local deviant). These sequences are presented repeatedly to the subject, forming a global regularity. A 5-tone sequence is considered “global standard” if it is identical to the preceding 5-tone sequences. Structure detection is then tested by measuring the subjects’ response to a global deviant sequence – i.e. a 5-tone sequence which is different from the preceding 5-tone sequences. If XXXXX is repeated multiple times, then the global deviant sequence is XXXXY, i.e a local deviant. Contrarily, if XXXXY is repeated multiple times, the global deviant sequence is XXXXX, i.e. a local standard. Therefore, this paradigm orthogonalizes local and global regularities. Experimentally, local deviants elicit a rapid mismatch negativity (MMN), whereas global deviants evoke a late (300 ms) electro-encephalogram potential (18).

### Structure Detection

We evaluated the model surprise to tones of a rand-regular sequence, where the last tone is changed into its binary opposite (Fig. 2 G). The model detected the simpler global deviant - local deviant (Fig. 2 H). Remarkably, it also detected the global deviant - local standard (Fig. 2 H).

### Learning dynamic

The global deviant - local deviant was detected after 3 weeks of pretraining, while it took around 3 month for the global deviant - local standard to be detected (Fig. 2 I, see supp. Fig. 4 for a zoom). We can’t make a one-to-one temporal mapping between these models dynamics and developing brains, as we present sounds without break and in batch of inputs. Nevertheless, it is remarkable that children at 3 month old are also able to learn such global deviance (42). 3 months is therefore an upper limit on the amount of data needed for this detection. Consequently the model learns with a reasonable amount of data, at least in order of magnitude.

### Effect of sequence complexity on structure detection

Since global deviants are harder to detect than local deviant, we wanted to investigate precisely how a structure’s complexity modulate its detection. Indeed the previous experiments do not distinguish if the model compress or purely store the sounds sequences, with an added cost for longer sequences. If the model were to compress the sequence, the structure detection would be faster for sequence which are simpler to compress independently of their size. Remarkably, humans do compress rather than purely store algebraic tones sequences. Indeed, Al Roumi et al. (16) observed that in humans, binary sequences with simple structure were memorized more easily than sequences with complex structure. More precisely, they proposed a set of 10 binary sequences, each composed of 16 tones (see methods for details on the complexity metric). Remarkably, humans detection performance was strongly correlated with the sequence complexity. We test the model on these algebraic patterns by embedding them in the rand-regular-rand protocol, as for repeating tones sequences.

### Structure detection

The model detected the structure of sequences of diverse complexity (Fig 2 K). Interestingly, the model surprise converged more slowly for the complex sequences: i.e. more tones are required for the model to change its surprise from the random baseline.

### Learning dynamic

The model’s ability to detect algebraic patterns emerged during pretraining (Fig 2 L). Remarkably, this emergence appeared in a sequential order, with sequences composed of chunks being accurately detected earlier (1-2 months, complexity <=6) than sequences composed of nested structures (3-4 months, complexity>=12) (see supp. Fig. 4 for a zoom).

### Controls

The model’s performances can be partially explained by the emergence of a trivial “copy” algorithm, where the model detects repetitions of an acoustic pattern instead of an actual algebraic structure. To verify that the model effectively represent sequences as algebraic structures, we tested its ability to generalize to novel sequences. In the regular part of these sequences, the 16 tones sequence is repeated 3 times with two novel distinct tones forming the sequence at each repetition. If the model detects the underlying structure, its surprise should progressively diminish with the number of repetition. In this “generalize” protocol, we also observed that the model could detect most structures and that this ability emerged through pretraining (supp. Fig. 5). The two most complex structures were not generalized by the model.

### Music but not speech pretraining accelerates emergence of regularity detection

The above results show that pretraining is essential to the emergence of structure detection. In practice, the models were pretrained with a variety of sounds: speech, environmental sounds and music. As these sounds have different statistical properties, it remains unclear which were used to support the emergence of structure detection. To address this question, we repeated our analyses on three sets of models pretrained solely on speech, environmental sounds and music, respectively. Two striking observations result from this analysis. First, exposition to speech alone did not allow the emergence of “zero-shot” structure detection (Fig. 3 A). Second, music exposure accelerated the emergence of structure discovery compared to environmental sounds (Fig. 3 B,C).

**Fig. 3.**
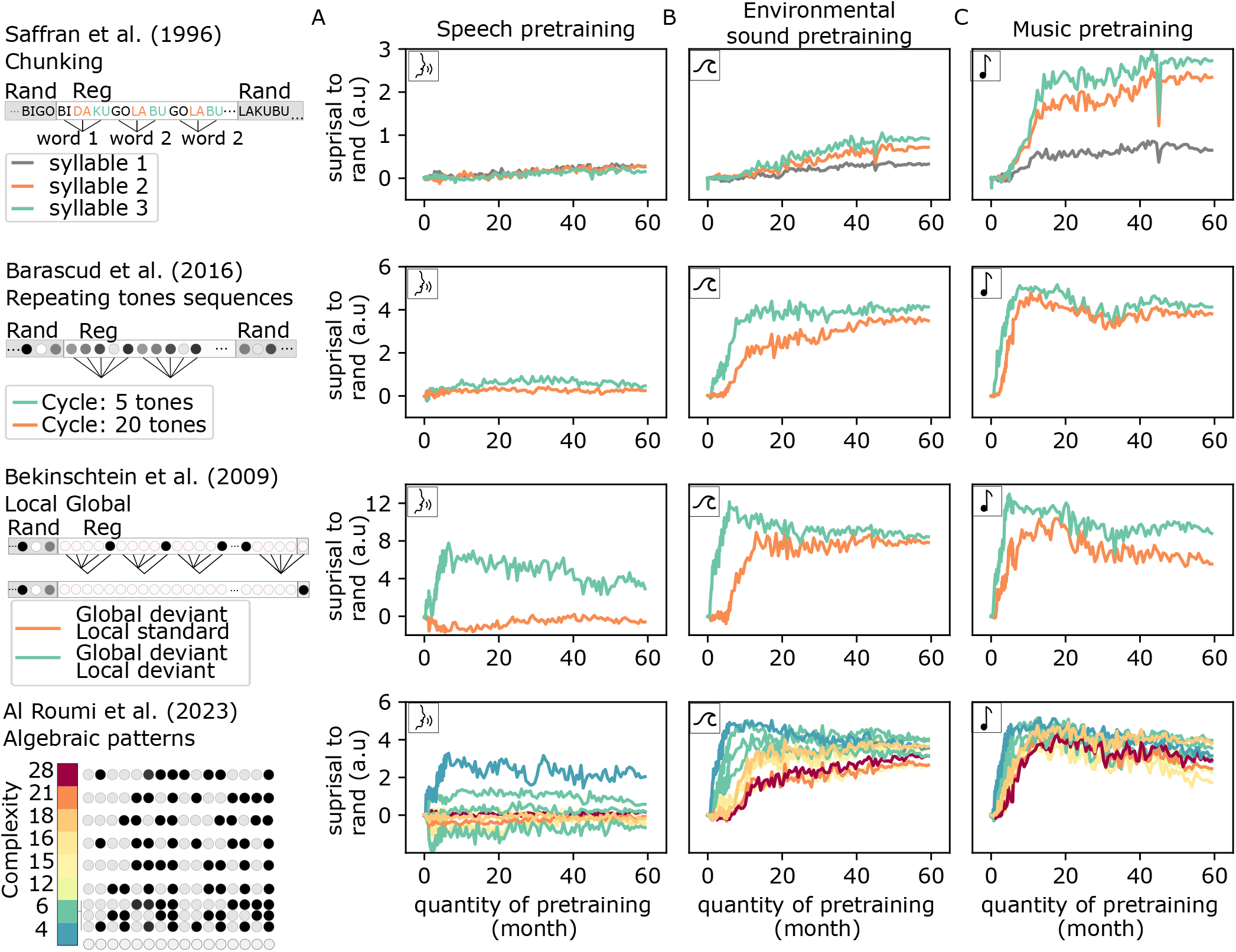
Effect of the pretraining dataset on the emergence of structure detection. We plot as a function of the amount of pretraining the ability of models trained on speech (A), environmental sounds (B) or music (C).

## Discussion

Overall, our analyses demonstrate that structure detection can emerge from self-supervised learning: A generic learning objective – unmasking natural sounds – allowed our neural networks to perform “in-context” structure detection, without any additional learning. Like humans, the spontaneous ability of a model to detect auditory structures depends on their algebraic complexity.

Together, these results challenge radically rationalist view, for which the neural representation of algebraic structures results from a built-in and specific mechanism. First, structure detection can be discovered through pretraining and thus needs not be an axiomatic property of the network (Fig. 2). Second, and to our surprise, this ability appears to be independent of speech processing (Fig. 3). While structures are undoubtedly critical for reasoning and syntax, our result highlights that the link between speech processing and the spontaneous detection of auditory structures may be more tenuous than previously assumed.

Two non-mutually exclusive interpretations may explain this dissociation between structure detection and speech abilities. First, our models pretrained with speech sounds may learn distinct types of regularities compared to those exposed to music: unlike music and many natural sounds, speech does not have many repetitions. This “asynchrony” of speech may thus lead the network to specifically discard repetitions and focus on other statistical relationships, like meaning, syntax and prosody. On the contrary, the “synchrony” of musical and of many natural sounds could favor the emergence of repetition detection, and, by proxy, the emergence of an efficient – algebraic – system of representations to compress these repetitions.

A second, alternative, interpretation could relate to an inherent weakness of Wav2vec2: unlike text-based models, such speech-based model appears to lack many high-level syntactic and semantic properties (e.g. (35, 47)). In contrast, the human brain may use additional algebraic representations to detect syntactic structures and recycle them to detect any auditory structure. This possibility, however, would need to reconcile the fact that mathematical, rather than language areas, seem to represent the algebraic auditory patterns (16, 48).

In this work, we systematically varied the type and the amount of auditory exposure used for pretraining. However, several other components remain to be tested. First, all of the presently-tested models used the same learning objective: unmasking. Yet, there exists other unsupervised objectives, such as causal predictions (49), compression (50), multi-losses and Hebbian learning. Second, we did not vary the architecture (36). In addition, Wav2vec2 presents several implausible features. We report in supplementary materials several weaknesses of the model. We observe that at large (>200ms) inter-stimuli interval, the model computation are biased by normalization operations. This normalization also leads the model to process differently sequences which differ minimally. This effects lead to surprising behavior, which can be resolved by the introduction of a few-shot protocol, introduced in the supplementary materials. More problematically, the architectures uses an auditory buffer which gives it a perfect short term memory over the presented sound waveform (e.g 20 s of sounds). On the contrary, it lacks any long-term memory module to recollect past episodes. These features are at odds with humans: For examples, humans can partially memorize random 20-tones sequences over several weeks (24, 51–53), presumably thanks to a dialogue between the hippocampus and the cortex (54–57). Overall, it is thus unclear whether our few-shot experiments do correspond to the memorization strategy effectively implemented in the brain. The fragility of the model behavior in the few-shot experiments, compared to the one-shot experiment, strengthens the idea that it misses modules for long term memorization. Finally, we showed that self supervised learning of naturalistic stimuli provide a sufficient basis for the detection of complex auditory structures. Yet, a core question remains: why then, are humans so much more efficient at such tasks than other animals? First, this cognitive advantage may not be that clear cut. For instance, ferrets do detect repeating sequences, but, unlike humans (20), this ability is limited to seven tones (58). Furthermore, while mice seem to fail to detect second-order violations (59), rhesus macaques and marmoset do detect such global structures (45, 46). Overall, we can thus speculate that, not only the amount of stimulus exposure, but also, and perhaps primarily, the sheer size of the cortex, may turn out to be a critical factor in the zero-shot detection of complex auditory structures.

Overall, the present minimal framework acts as a proof of existence: algebraic structures can spontaneously emerge in generic neural networks trained on natural sounds with self-supervised learning. Perhaps more importantly, the quick development of efficient models and learning rules pave the way to test, empirically, how built-in and acquired mechanisms may contribute to humans’ unique ability to represents complex structures.

## ACKNOWLEDGEMENTS

The author would like to thank Stéphane Deny, Fosca Al Roumi, Christophe Pallier, Linnea Evanson, Théo Desbordes, Sam Norman-Haignere and Stanislas Dehaene for comments on the work and manuscripts.

This work was granted access to the HPC resources of IDRIS under the allocation 2023-AD011014524 and 2022-AD011013176R1 made by GENCI (P.Orhan). The author would like to thanks IDRIS for their support on the Jean-Zay cluster. This project has received funding from the European Union’s Horizon 2020 research and innovation program under the Marie Sklodowska-Curie grant agreement No 945304 (P.Orhan). This work was funded in part by FrontCog grant ANR-17-EURE-0017 (Y. Boubenec and JR. King for their work at PSL).

## Methods

### Models pre-training

Models are pretrained with Huggingface’s distributed trainer (60), on a set of 64 V100 GPUs (8 gpus per node) for 100 000 steps of gradient descent, with a batch of size 4 in each GPU (each training took approximately 4 days). The training follows all recommendations and hyper-parameters used to train the base Wav2vec2 model in (36). Notably, we make several modification to the implementation of (60) since it missed some important elements of (36). First, we scale the gradient of the encoding features by a factor 10, which has for effect to slow down the learning of the encoder compare to the transformer part of the architecture. Then, we multiply the mean of the squared encoding features (the features at the output of the convolution) by a factor 10 and add this value to the total loss as a regularizer. If we didn’t have these scaling, the model activity would either diverge or collapse during training. We train with input sounds truncated to have a maximal size of 20s and minimal size of 2s. For each dataset, we pretrain two models from different random initialization and random seed throughout pretraining. The results were all consistent across the pairs of models, demonstrating that the difference of results did not arise by chance. Moreover, the losses of the models during pretraining were similar across the 8 trained networks. To make sure our models had converged to reasonable solutions, we tested their classification performances on speech ( classification of logatomes), music (classification of musical genres), and environmental sounds (classification of environmental scene). More precisely, for the speech test, we took a subset of the OLLO2 dataset (61). The model is trained to predict a set of logatomes identity (9 distinct logatomes) from a subset of speakers and we test its ability to generalize across speaker. More precisely, we learn a cross-validated data normalization and logistic regression from the model time-averaged response to each sound and the class label. Scores are collected across cross-validation folds and across layers of the model and averaged. A similar approach is used for the music test with the GTZAN dataset (62), where the model needs to classify 30 distinct musical genres. We make sure to use error-free cross-validated folds (63). Last, we use the ESC-dataset (64) for classfication of environmental scenes. We here test the unsupervised model representations at the end of pre-training (no fine-tuning). Remarkably, models pretrained on a specific type of data performed better on the respective test than models pretrained on a distinct type of data (supp. Fig. 7). Additionally, models trained on everything performed well on all the test. These observations confirm that models have converged to reasonable solutions and are able to perform the contextual processing required for these classifications.

All models weights of all checkpoints and the exact datasets used for pretraining are available upon reasonable requests to the author.

### Sequences generation

Syllables were generated using the MRBOLA synthesizer, through the python package voxpopuli ^3^. We used the english voice with a speed of 160. Each synthesized syllable was generated to be of duration 200 ms, and re-sampled at 16000 Hz and normalized to have a root-mean square of one. We also removed any pitch-modifier from each syllable during synthesis.

Tones were generated using librosa (65) and lasted 50 ms. Both tones and syllables were multiplied at their onset and offset with hanning windows (i.e raised cosine window) of size 5 ms to avoid artefactual clipping. For figures 2 and 3 we used sequences without any silences between tones or syllables. For, and only for, supplementary figure 8 we removed the waveform normalization. With waveform normalization, the model is not robust to repeated silences and it became unable to detect structure at ICI larger than 0.2s. Without waveform normalization, the model is still not robust to silences but its performances decreased at a larger ISI.

### Loss measurements

Evaluating correctly the loss a masked network like Wav2vec2 requires some careful modifications to its original code base and careful experimental design. We have not seen earlier description in the literature of these points of attention. To facilitate further research, we describe in details the steps taken and provide our implementation of the needed modifications.

Wav2vec2 is equipped with a contrastive loss, as such it requires the selection of a set of masked latent vectors and for each of these latents a corresponding number of latents distractor from the same acoustic context, termed negatives. Each latent is sampled every 320 time-step i.e every 20 ms (50hz), and gather information across a receptive field of 400 time-step.

To prevent leakage of information when computing Wav2vec2 loss, it is extremely important to mask all latents whose receptive field overlaps with the presence of a certain sound. When computing the loss for a certain sound event, we therefore masked all latents whose receptive field overlapped with this event. We compute a loss value for all these masked elements, but kept and summed only the ones whose receptive field was fully contained in the sound event, producing a final loss value for this sound event.

Let us go through an example: given a 10 second waveform *w ∈* R^160000^ discretized at 16000 Hz, the model computes a latent speech representation of dimension 512 *q ∈* **R**^(512,499)^ at 50 Hz (the exact number of latent is 499 here, and can be computed from the strides and kernel sizes of the successive convolutional layers). These latents are separated by 20 ms, but they are the result of successive convolutions, such that their value is influenced by initial waveform values of up to 25 ms in the future. Consequently to measure the surprise of a tone happening at 2s and lasting for 50 ms (2 to 2.05) it is not sufficient to mask the 100-th latent ( receptive field 2 to 2.025) . We need to mask the latent 99 (receptive filed 1.98 to 2.005), 101 (receptive field 2.02 to 2.045) and 102 (receptive filed 2.04 to 2.065), but not the latent 98 (1.96 to 1.985) or 103 (2.06 to 2.085). Additionally, one needs to set to 0 all sound latent happening after 102, such that the relative positional embedding present in Wav2vec2 does not also introduce leaking of future information.

The contrastive loss contrasts the prediction over the masked latent with its true unmasked value and a set of other unmasked latent values, termed negatives. To obtain a reasonable values of the loss, the set of negatives should be shared across masked latents, solely removing the negative that match the masked latent. Otherwise, the loss might purely reflect the acoustic distance between the embedding of the acoustic events composing the set of negatives. In that case it would poorly reveal the network errors in predicting, correctly or not, the masked elements. To prevent such problems, our experimental design make sure to have sufficient negatives by using a random sequences composed of all the tones or syllables from the alphabet at the beginning of the sound. This design is therefore due to a limitation from the way negatives are sampled in Wav2vec2.

The model is evaluated in a partially causal way. More precisely, to measure the response to each tone, we set to 0 the activity of all future latent, critically before the positional embedding is computed. Note that it only makes the model partially causal because we do not: (1) set the attention masks used by the transformer to be causal; (2) adapt the positional encoding to take into account this causality. The reason is that Wav2vec2 is pretrained in an acausal regime, therefore it showed poor performance when evaluated with causal attention masks, which is the standard way to make a model causal. Nevertheless, the loss we measure remain only influenced by past sounds and is consequently a causal measure. Future work will adapt and pretrain novel models to solve these issue. These models will be able to perform in a single pass all causal predictions over a sound.

We evaluate the model over 371 checkpoints across the 100 000 training steps. More precisely we save the checkpoints from step 1 to 100 by step of 1, from 100 to 1000 by step of 10, from 1000 to 10 000 by step of 100 and from 10 000 to 100 000 by step of 1000. This schedule allowed to have a precise overview of the beginning of training where most of the interesting dynamics occur, while leaving the possibility to observe long term dynamics.

We would like to note that there are two limitations concerning the probing of the model contrastive loss. First the model has to be probed as many times as there are sound elements. Second, a decision of putting a mask over certain tones is explicitly made, which acts as a prior that guides the model behavior. Future architectural work will consequently resolve these two issues. For the first, by developing transformer model allowing parallel query of the loss for a set of sound elements partitioning a sound. For the second, by developing internal models that predicts the mask to be used in a particular context. The first development is a matter of architectural tweak, slowed down by hardware bottlenecks. The second will require more research, for example with the use of oscillatory mechanisms to guide masking. This matter is pressing, as loss probing with masked auto-encoder is computationally very inefficient. The probing of the networks activity and loss required around 30 000 hours of GPU compute time during the project, which could not have been done without a cluster. We estimate the carbon footprint of the project to be around 0.6 tones of CO2, based on the grid consumption value provided in (66).

### Complexity metric

We use the complexity measure investigated in (67) and (16). The sequence complexity was measured as the minimal length needed to write the sequence in a certain language, termed binary Language of Thoughts (LOT). We use the software from (67) to evaluate the LOT complexity of our novel sequences.

## Supplementary Note 1: Supplementary results

**Supplementary Figure 1.**
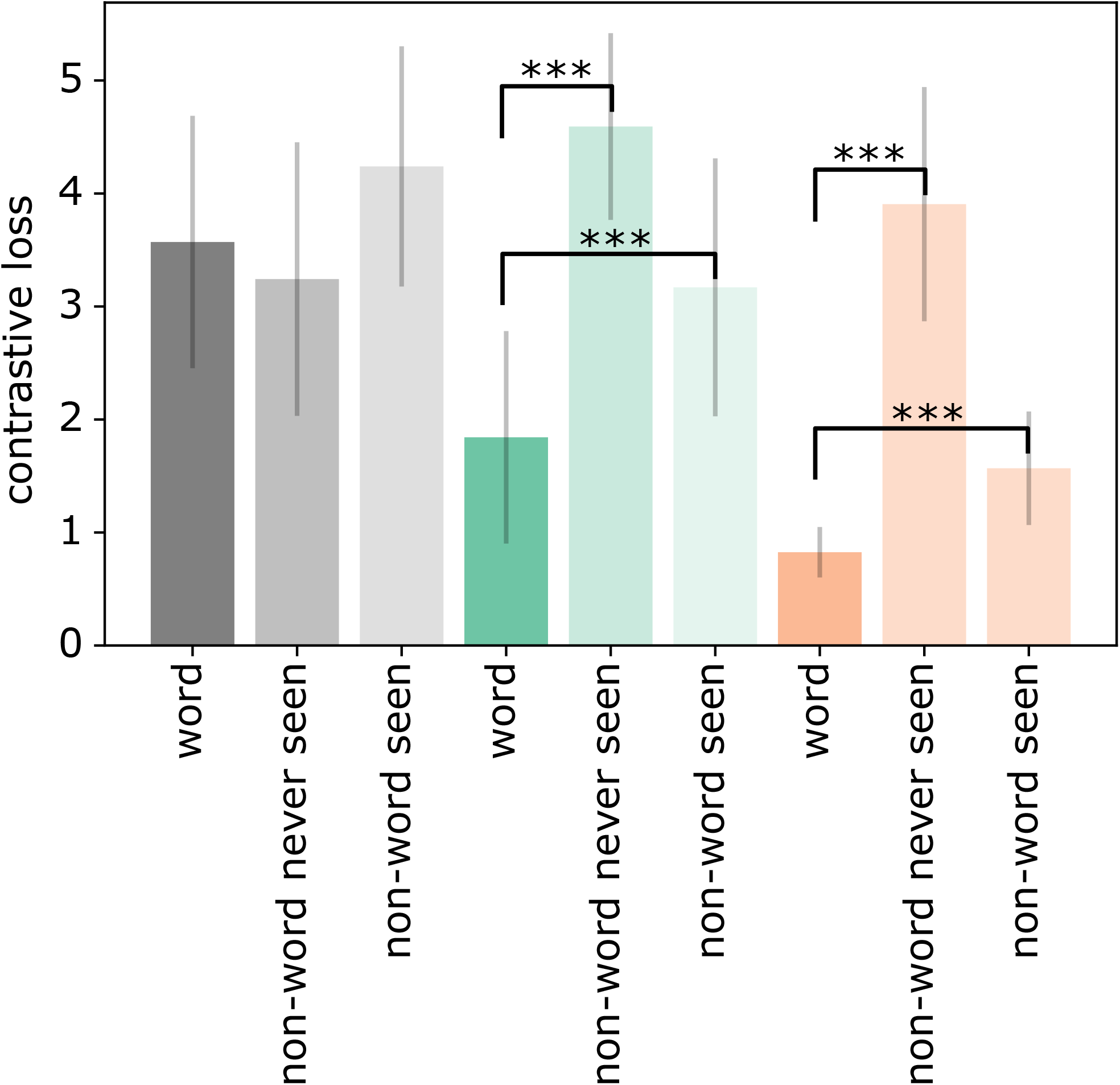
Contrastive loss to seen word, non-word never seen and non-word seen. Left three bars plots the average contrastive loss for the first syllable, middle second and right the third. Error bars capture the standard deviation across 30 trials. If the word is detected, the surprise, i.e contrastive loss, should be smaller at syllable 2 and 3 for the non-word seen and even smaller for the non-word never seen. This is indeed the case for syllables 2 and 3 (t-tests, p-value<0.05)

**Supplementary Figure 2.**
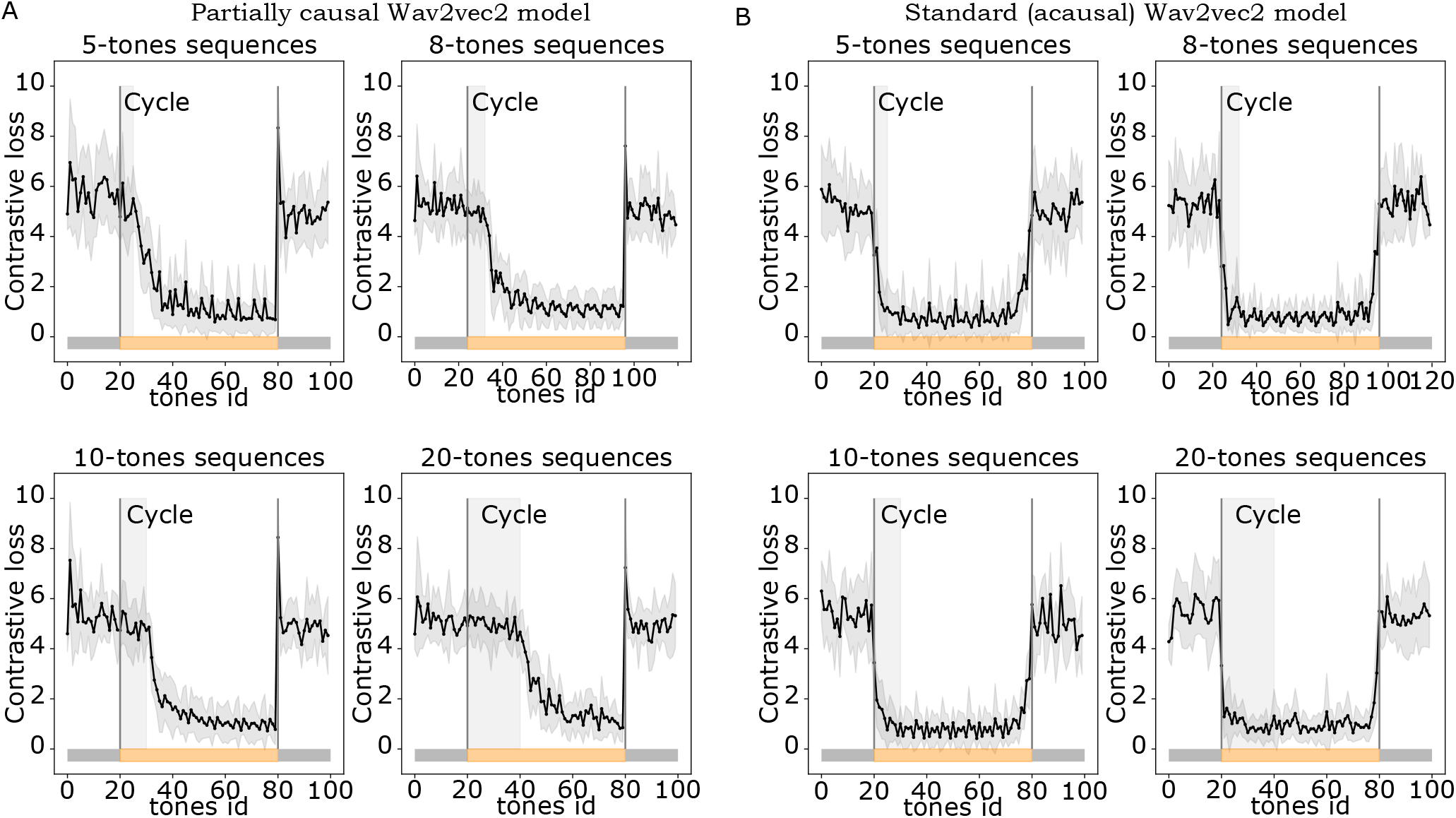
Contrastive loss of a causal (left) and acausal (right) models in responses to N-tones sequences (*N ∈ {*5, 8, 10, 20*}*). The acausal model detects the regularity earlier than the causal model.

**Supplementary Figure 3.**
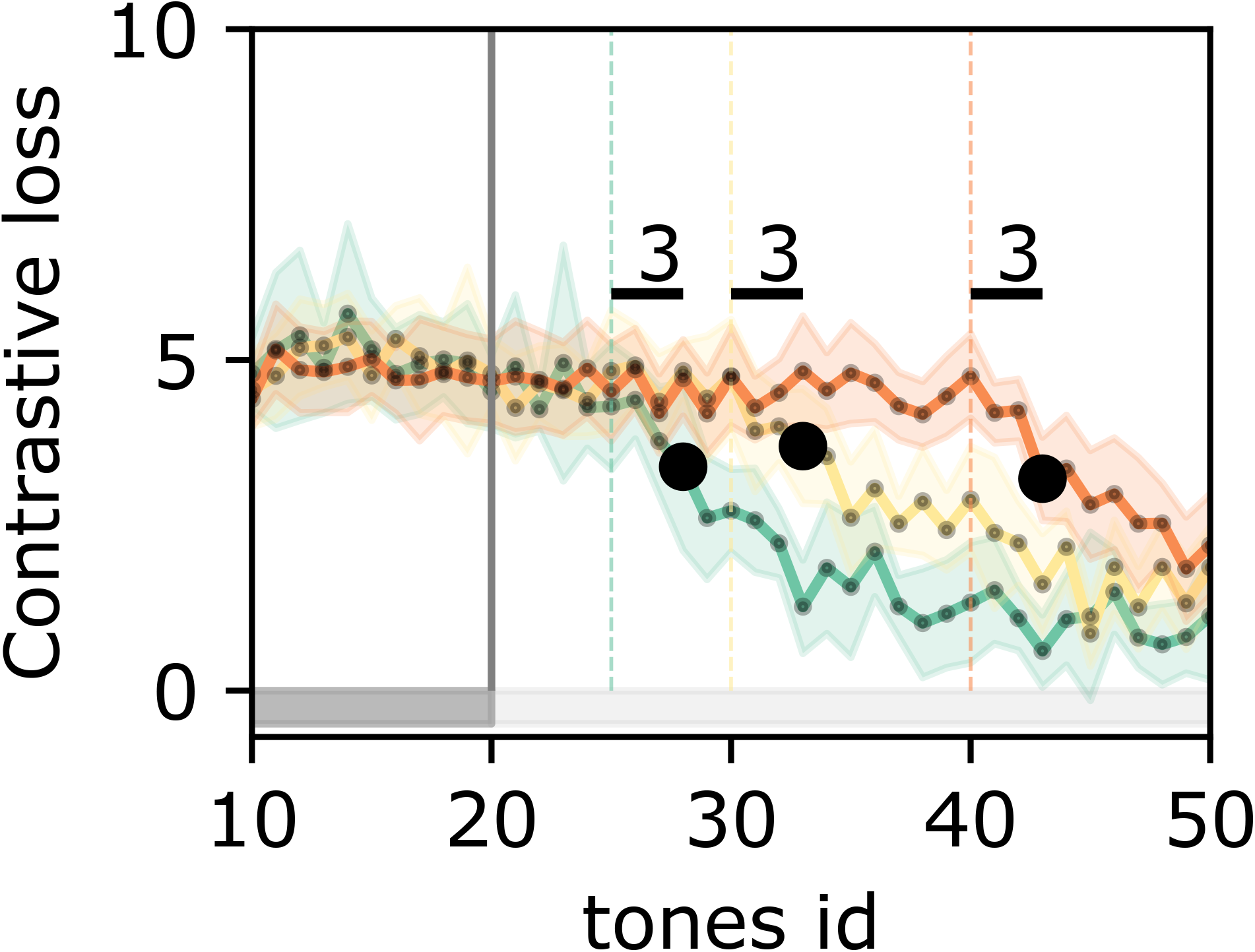
Zoom on the transition from random to regular. We mark as black dot the first tones whose contrastive loss were significantly different from the contrastive loss of the final set of random tones. We indicate in black the distance between this tone and the end of the first cycle. The significance is assessed with t-test for independent samples across the 30 trials, and we choose for p-value threshold 0.005. Note that this result is dependent on the p-value threshold chosen, such that we would not qualify it as a robust results. Nevertheless, the models is in the order of magnitude of the number of tones at which the detection should happen as predicted by an optimal bayesian observer in (20), and at which humans detect the tones.

**Supplementary Figure 4.**
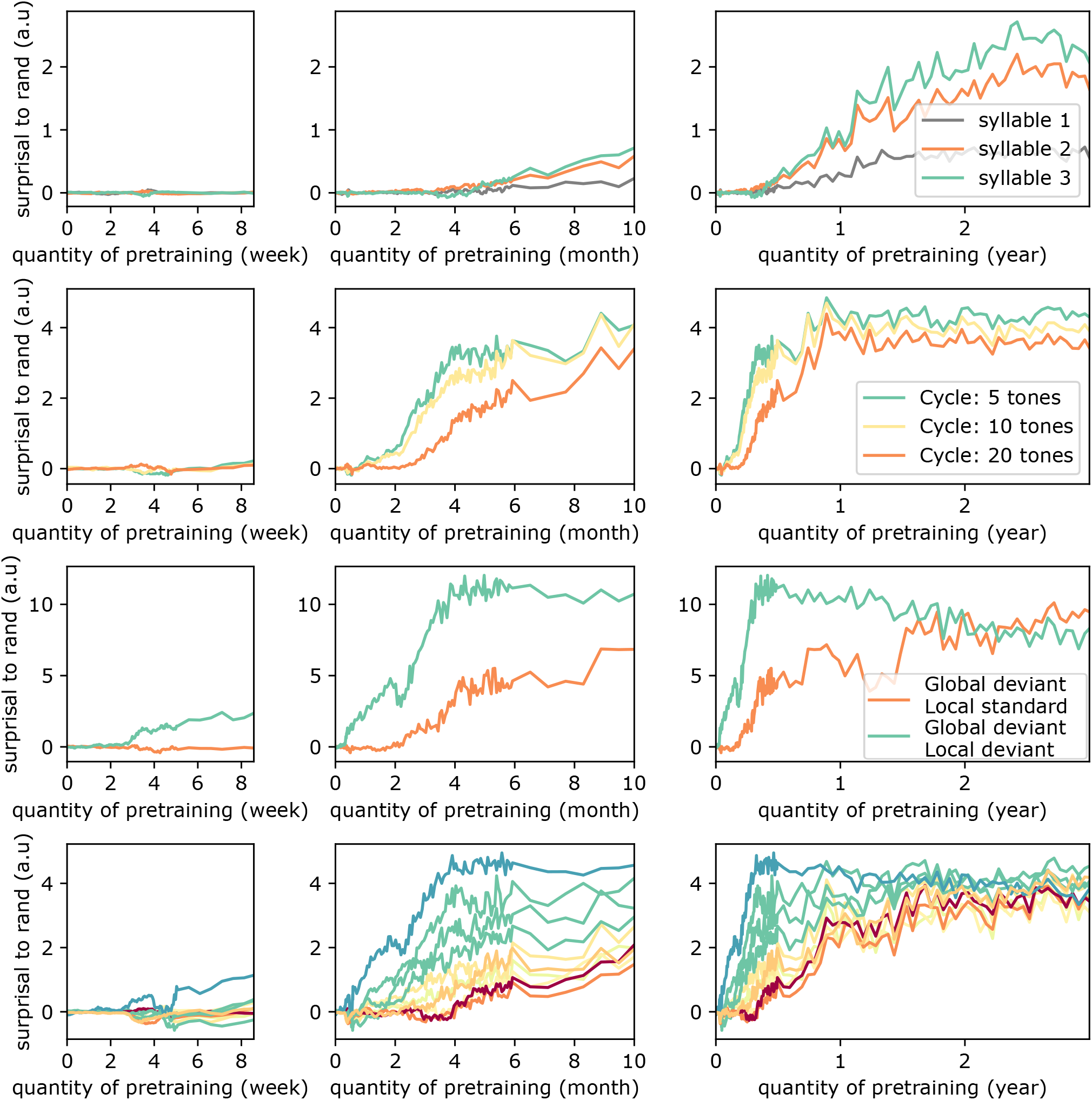
Zoom on figure 2, for each of the four experiments. First row: note the late emergence of syllable chunking, starting between 6 and 8 months. Second row: note the 2-months delayed emergence for the longer 20 cycle sequences compared to sequences of cycle 5 and 10. Third row: note the later emergence of Global deviant - Local standard detection compared to Global deviant - Local deviant. Fourth row: note the sequential emergence as a function of the sequences’ complexity between 2 and 6 months.

**Supplementary Figure 5.**
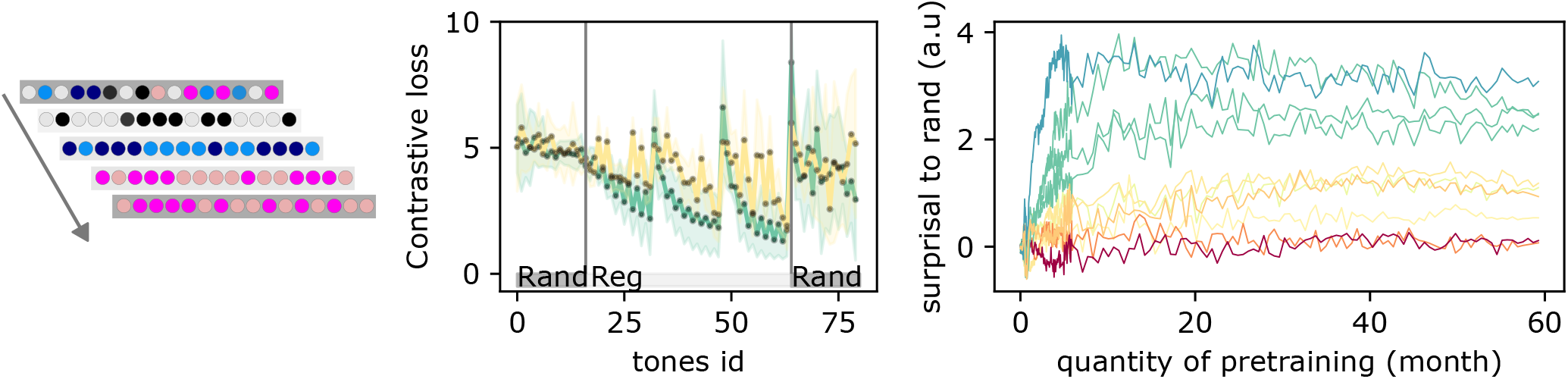
Generalization protocol. We test if the networks generalizes to sequences with novel sound elements but shared structure. Sequences are the same as before, i.e, the algebraic patterns of (16). At the onset of each repetition of the structure, the loss peaks because the model doesn’t know the novel elements. In the third and second repetition, as the model detects that these novel elements follow the same structure, the loss quickly decreases, to finally reach a lower surprise than the preceding sequences, and demonstrate a strong surprise to a random novel sequence. We measure on the right, the difference between the loss over the last 10 tones of the third sequences and the random pattern. Indeed we can’t use the loss over the last 16 tones, as we need to let the model see at least 2 tones, which happens at the 5-th tones for some sequences, forcing the use of the last 10 tones only.

**Supplementary Figure 6.**
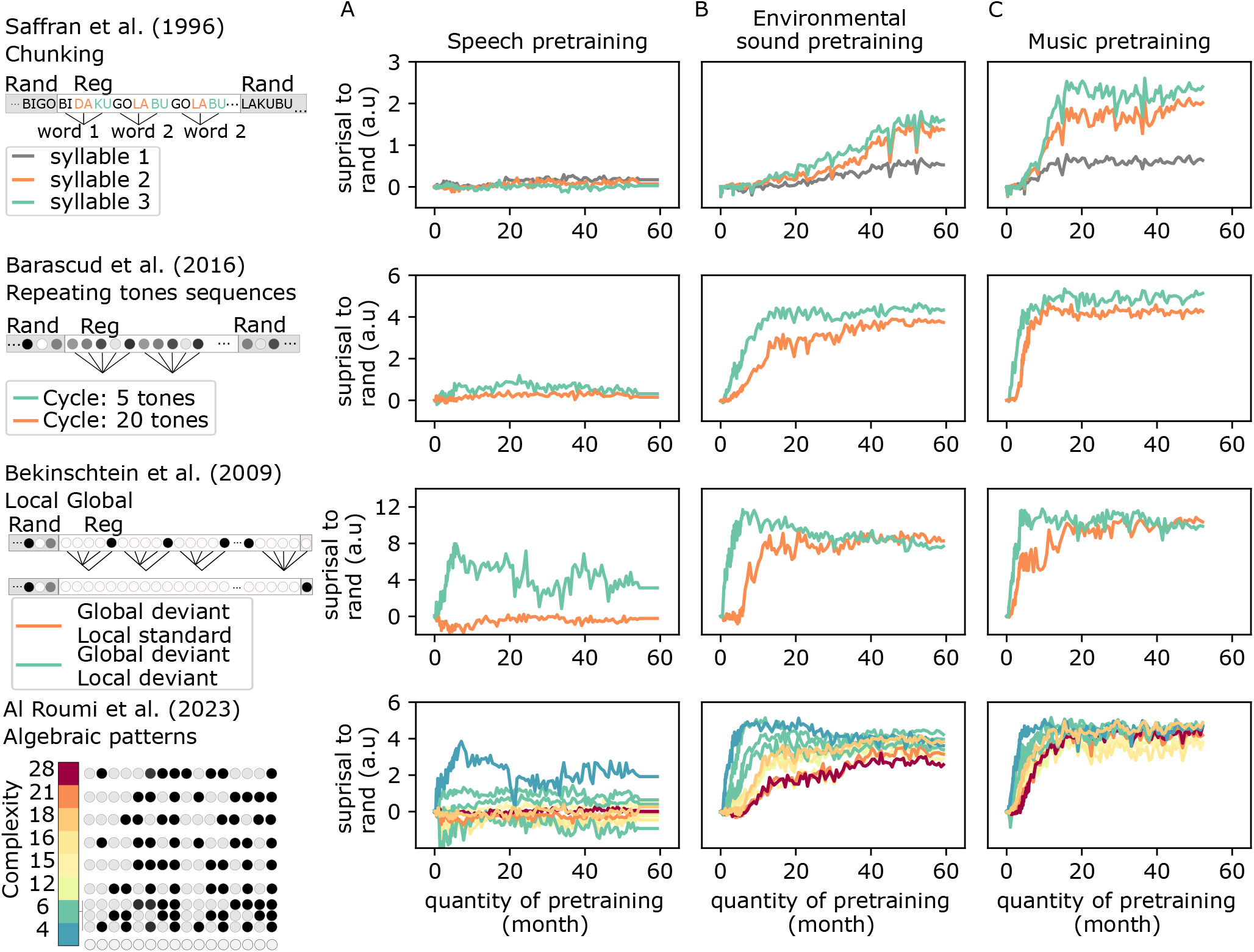
Same as figure 3, but evaluated on a second set of pretrained models. We observe the same dynamics as in the original figure, indicating that our results are robust to the random initialization of the models before pretraining.

**Supplementary Figure 7.**
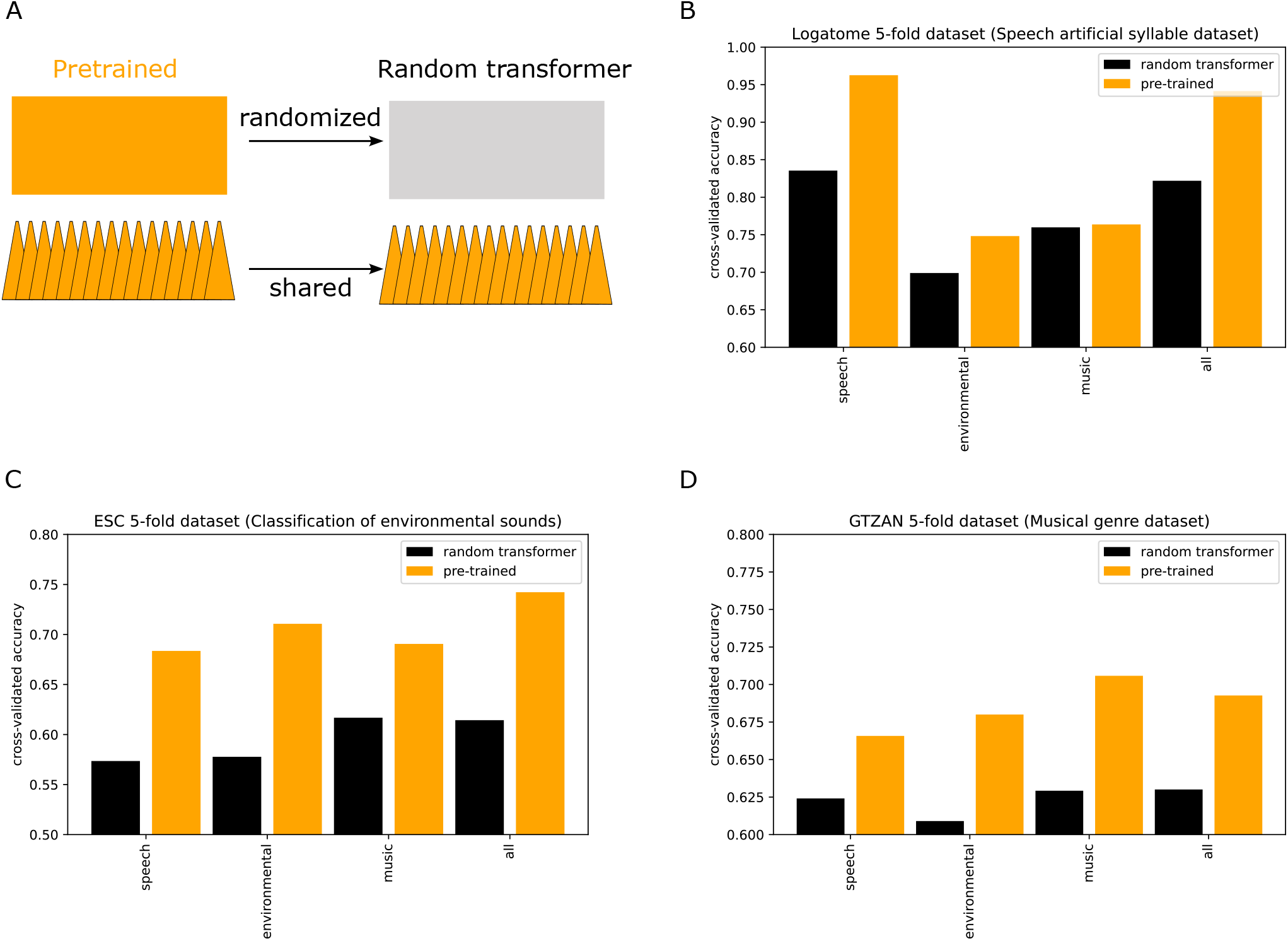
A. We evaluate the pretrained models’ representations in two conditions. Models were either unchanged (left) or their transformer layers randomized while preserving the learned parameters of their convolutions layer (right). B: Classification performances on logatomes. C: Classification performances on an environmental sounds. D: Classification performances on musical genres. Comparing orange and black bars allow to perceive the added benefit of contextual integration versus the purely local filtering of the convolutions. The black bars is not to be taken as a fully random baseline, which would require randomizing the convolutions as well.

**Supplementary Figure 8.**
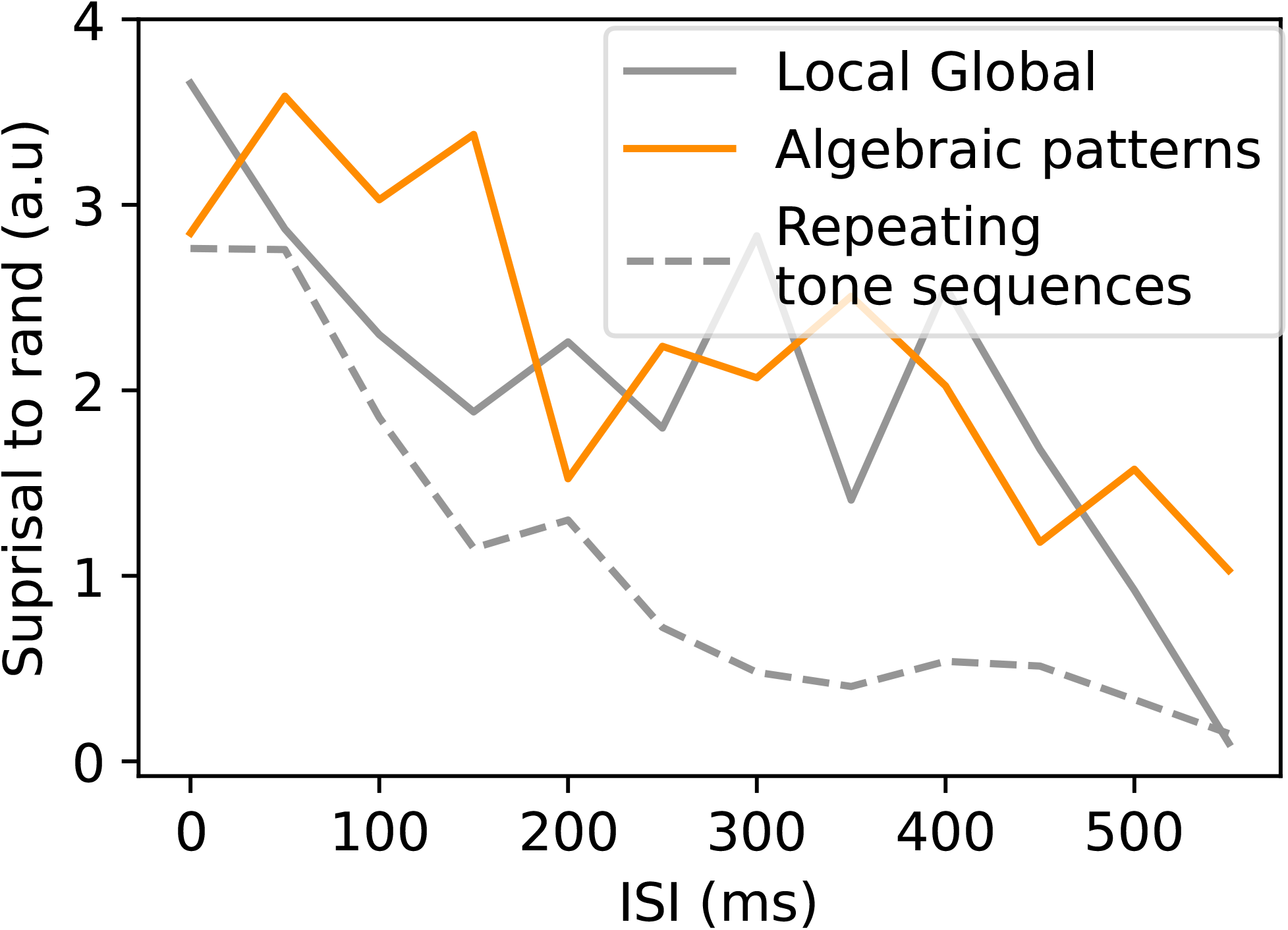
Decrease of the one-shot detection ability at large inter-stimuli interval (ISI). We measured the loss gap between random and regular sequence as a function of the silence duration between tones used for the binary-tones and N-tones experiments.

### Structure with longer inter-stimulus intervals (ISI) are harder to detect

We questioned if the model was robust to the insertion of silences in the sounds. We therefore added a constant inter-stimuli silence interval (ISI) between elements of the repeating tone sequences and algebraic sequences. We then repeated our evaluation with the model trained on all sound types. Problematically, we observed that the model started to diverge from its expected behavior at large ISI. For example, in the presence of silence, the model’s surprise to a repeating tone increased with the number of repetitions, whereas it should stay low near 0. We hypothesized that this effect was due to the normalization occurring on the waveform and after the first layer of convolution. Indeed, in the presence of a repeated element or repeated silences, this normalization biases the extracted local representation, which in turn impairs the contextual processing. Removing the waveform normalization helped mitigate this issue but the problem remained at longer ISI due to the normalization in the second convolutional layer (Supp. Fig. 8). Removing or changing this normalization to be computed per non-silenced sound bouts did not improve the model behavior. This issue will therefore have to be resolved by a change in the model architecture and novel pretraining.

Nevertheless, in ISI ranges typically used in experiments tackled here (200ms for (16), 150ms for (18)), the model has a reasonable behavior (Supp. Fig. 8). The model performed robustly to additional silences of 0.5 or 0.75 seconds between algebraic sequences. For repeating tone sequences, the slight decrease of performance with silence of 200ms mirrored the recent results of (68), where participants performed the same task as in Barascud et al. (20) but with increased difficulty as the silence gap increased.

Overall, we conclude that the model lacks robustness to silences, especially because of two successive normalization. Removing the first operation of normalization helps, and in this regime, the model detects structures at experimental ISI ranges. In the regime of over-normalization, the model abilities collapse for silences beyond 200ms. We therefore conduct further explorations, reported in details bellow, questioning if the model abilities can resurface through a few-shot scenario. In this scenario the model is allowed to learn from backpropagation and tested on deviant sequences. The model ability were recovered after a small (<50) number of backpropagation steps. We therefore hypothesize that further improvement of the models, for example by forcing silences to be present during pretraining, could be sufficient for the model to be robust at all ISI.

**Supplementary Figure 9.**
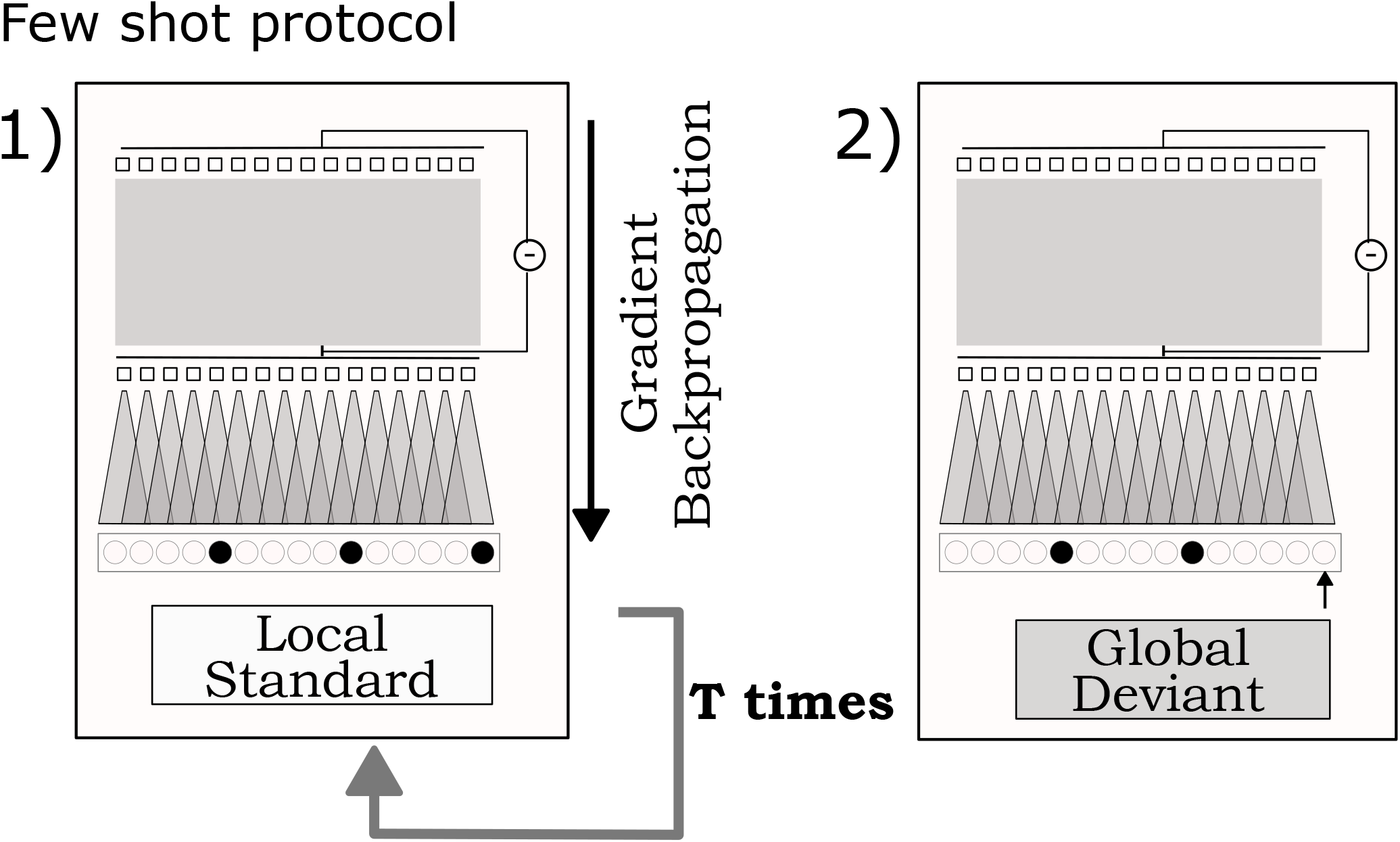
Few-shot protocol. 1) A sequence (here the Local Standard of the Local Global paradigm) is presented repeatedly to the model. Between each presentation, the model parameters are updated through backpropagation. 2) The model surprise on a deviant sequence is then measured (here the Global Deviant of the Local Global paradigm).

### Learning from few-short exposure to regular structures

Can online self-supervised learning improve the detection of repeated structures? In all of the above experiments, the model was evaluated in “zero-shot”, meaning that test stimuli did not change any of the internal parameters of the model. While zero-shot detection cannot trivially depend on synaptic plasticity, repeated exposure to the test sequences unfolds over long time periods that allow synaptic plasticity to change the network. Therefore, we tested a more permissive – yet plausible – “few-shot” scenario, where the model was plastic and could change its internal parameters upon successive presentations of the repeated structure. Specifically, we mirrored human experiments (16, 20), and repeated the standard sequences over several seconds to minutes. The model could adapt to the test stimuli through several gradient descent steps (Supp. Fig 9), using the same unmasking objective used for pretraining. We tested whether this few-shot exposure led to enhanced extraction of standard sequences and algebraic structures by evaluating the contrastive loss on the deviant sequences. To face the model with challenging stimuli, we used ISIs of 250 ms. To track the learning of the models over stimulus repetitions, we tested the model every 5 checkpoints from step 5 to 50, and then every 50 steps up to 600.

### Repeating tone sequences

We first tested the effect of few-shot exposure with repeating tone sequences at an ISI of 250 ms. The habituation was composed of four repetitions of the tone motif. In the test sequence, the fourth sequence was changed by swapping one tone for a deviant tone (Supp. Fig. 10 A). The model detected repeated structures in the zero-shot scenario, but improved with repetitions of the standard sequence (Fig 10 B).

### Global deviant - local standard

Next, we habituated the model to the XXXXY sequence and tested its ability to be surprised by a global deviant - local standard token in last position (XXXXX) (Supp. Fig. 10 C), with an ISI of 250 ms. Habituation sequences were composed of three repetitions of the local deviant sequence: XXXXY. The test sequence was composed of two repetitions of XXXXY followed by the global deviant - local standard XXXXX. The ability to be surprised by the global deviant - local standard emerged (Supp. Fig. 10 D) but only when the model was exposed at least 35 times to the sequence. We repeated the experiments for different learning rate parameters and across the eight fully-pretrained models, but none were able to detect the global deviant in less than 20 steps of gradient descent (Supp. Fig. 11).

### Algebraic patterns

We tested the model ability to memorize algebraic patterns by measuring its sensitivity to 4 alternative deviants per sequence, as used in (16) (Supp. Fig 10 E). The model was able to memorize all sequences, but simpler sequences were memorized faster than more complex sequences (Fig Supp. 10 F). As humans, after being exposed to 10 repetitions, the model sensitivity was correlated with sequence complexity (R = *−* 0.86 *±* 0.05, statistics across the last 10 pretraining checkpoints). We performed several controls to further explore this finding. First, the memorization of each sequence could depend on their duration, which largely differ (Figure 1A). Sequences with smaller cycle are more repeated and therefore could be learned faster, which would explain the correlation between sequence complexity and learning speed. To test this hypothesis, we generated 10 novel sequences which were all of duration 16 tokens, but varied complexity. Model sensitivity stayed high (min D’=1.57, max D’=4.0, mean D’=3.56) and still correlated with sequence complexity (R = -0.66, Supp. Fig. 12). Second, we tested if the model could generalize structure detection to novel sequences, based on the same algebraic pattern but different sound elements. We replaced the repeated exposure to standards with the repeated exposure to 3 novel sequences, each composed of different sound elements while preserving a shared algebraic pattern. We tested the model on a fourth sequence, also composed of novel sound elements but the same shared algebraic pattern. The model generalized from all but the most complex pattern, with other nested patterns requiring at least 50 repeated exposure (Supp. Fig. 13). Finally, we stress that despite the model ability to perform few-shot detection at long ISI, the model behavior and dynamic of emergence are much noisier in this few-shot scenario than during the one-shot scenario at low ISI (Supp. Fig. 11, 12,13). Therefore, such models have to be improved to exhibit structure memorization as robustly as humans.

### Methods for the few-shot protocol

To test if the network is able to learn sequence structure if we turned on plasticity, we pursued its pre-training by performing 600 gradient descent step using the exposed sound. As for pre-training, we used the HugginFace Trainer implementation (60), with the same modification to the Wav2vec2 implementation of HuggingFace as for pre-training (replicating the original Wav2vec2 implementation available in FairSeq). The Adam optimizer used during pre-training implements a learning rate scheduler, leading to small weights modifications at the end of pre-training. To maximize the effects of the exposition, we therefore re-initialize the optimizer for each of these test. The ADAM optimizer learning rate decreased with a linear ramping from a maximal value to 0 across these 600 gradient descent steps. We explore several learning rate maximal value: 2.5 *∗ ×* 0^*−*4^, 0.75 *×* 10^*−*4^, 2.5 *×* 10^*−*5^, 0.75 *×* 10^*−*6^, 2.5 *×* 10^*−*6^. Results were consistent across learning rate, except at the largest one where no convergence was observed. We report results for a learning rate of 2.5 × 10^*−*5^ in the main paper, and results for all learning rate can be observed in the supplementary.

Across steps, we first iterate through the different masks, with each mask covering one of the sound element present in the sequence, and then through 3 repetition of the same sequence (detection set-up) or through concatenation of 3 sequences with shared structure but different tones. These 3 sequences mirror the “zero-shot” scenario where they were concatenated one after the other. At test time, we use either the same sequence (detect) or a novel sequence with the same structure but novel sound elements (generalize). The optimization is performed with batch of size 16 and uses a single GPU. A batch of size 16 allowed to perform gradient descent simultaneously through all the masks corresponding to the 16 tones of the binary tone sequences. During these gradient descent steps, the model employs the same regularization tool as during pre-training, notably unit dropout and layer dropout. We observed that a dropout of the first layer generated loss spikes, which disappeared when layer dropout was not allowed. Although this could participate in the fragility of the model in the few-shot scenario, we preserved this dropout to mirror in few-shot, the updates done in pretraining, and, more generally, stick to the hyperparameters found by the original author of the Wav2vec2 model (36).

Finally, note that the HuggingFace Trainer uses a gradient scaler for technical purposes. We observed that this scaler lead to the skipping of the first few step of optimisation to find an adequate gradient scaling factor, notably because we re-initialized the optimizer. We recorded which of those steps were missing, and accounted for them such that step five reported here was exactly the fifth step of gradient descent. These variations have therefore no impact on the results we report here.

### D’ measurement

The D’ is estimated from the model loss to all sound elements of four different deviant sequences. For each of these deviants sequences, a different sound element is replaced with another from the sequence. For example, one tone is replaced with its binary alternative in binary sequences. For a sequence of size N, a threshold on the loss then determines which elements are classified as predicted or not-predicted by the model. To have high sensitivity, every elements among the N elements of the sequence should be classified as predicted, except the deviant which should not be well predicted (i.e deviants should have a larger loss than other elements). The threshold determines which elements are predicted versus non-predicted. We compute the sensitivity of this particular classification by merging the results across the 4 deviants sequences and computing a D’ from this set of 4N classifications. This operation is repeated for all of the 4N possible threshold values. More precisely, we change the classification boundary so that one element changes class at a time, starting from the one with the largest contrastive loss to the one with the smallest contrastive loss. We report the D’ measure of the model sensitivity on this sequence as the average of these 4N measures, weighted by the 4N loss gaps between each threshold change. This weighting procedure allows us to take into account how far the loss was on the deviants compared to other elements. We observed manually that this measurement captured well the model’s behavior.

**Supplementary Figure 10.**
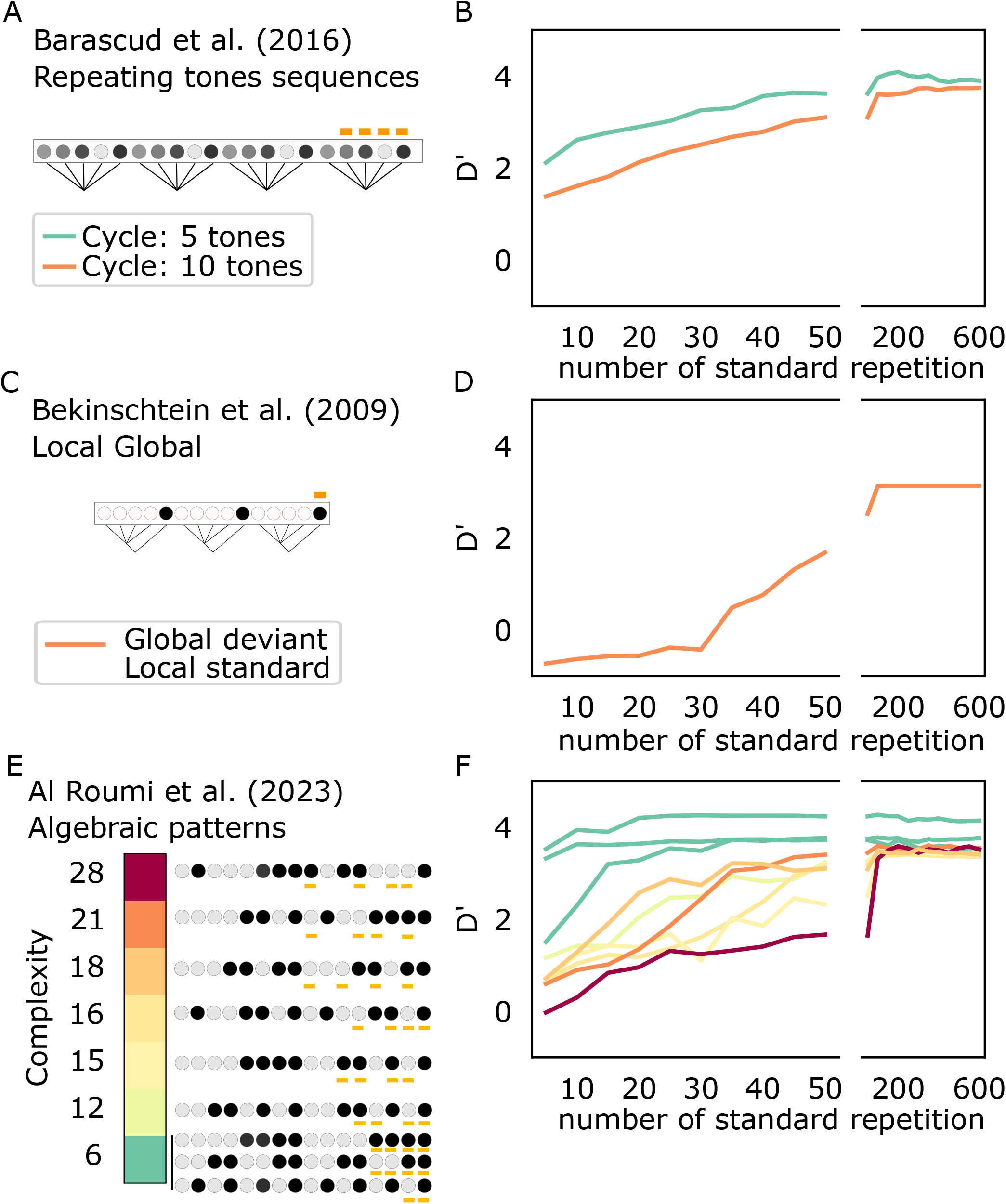
Self-supervised gradient descent led to sequence memorization modulated by structure complexity. A: The model was exposed repeatedly to four repetitions of a random sequence of 5 or 10 tones, with 250 ms inter-tones interval. The model was then tested on deviant sequences, in which one tone (marked by orange ticks) was swapped for another deviant tone. B: Model sensitivity to deviants as a function of the number of exposition. C-D: Same as A-B but for the local global paradigm of (18). E-F: Same as A-B but for algebraic patterns of (16).

**Supplementary Figure 11.**
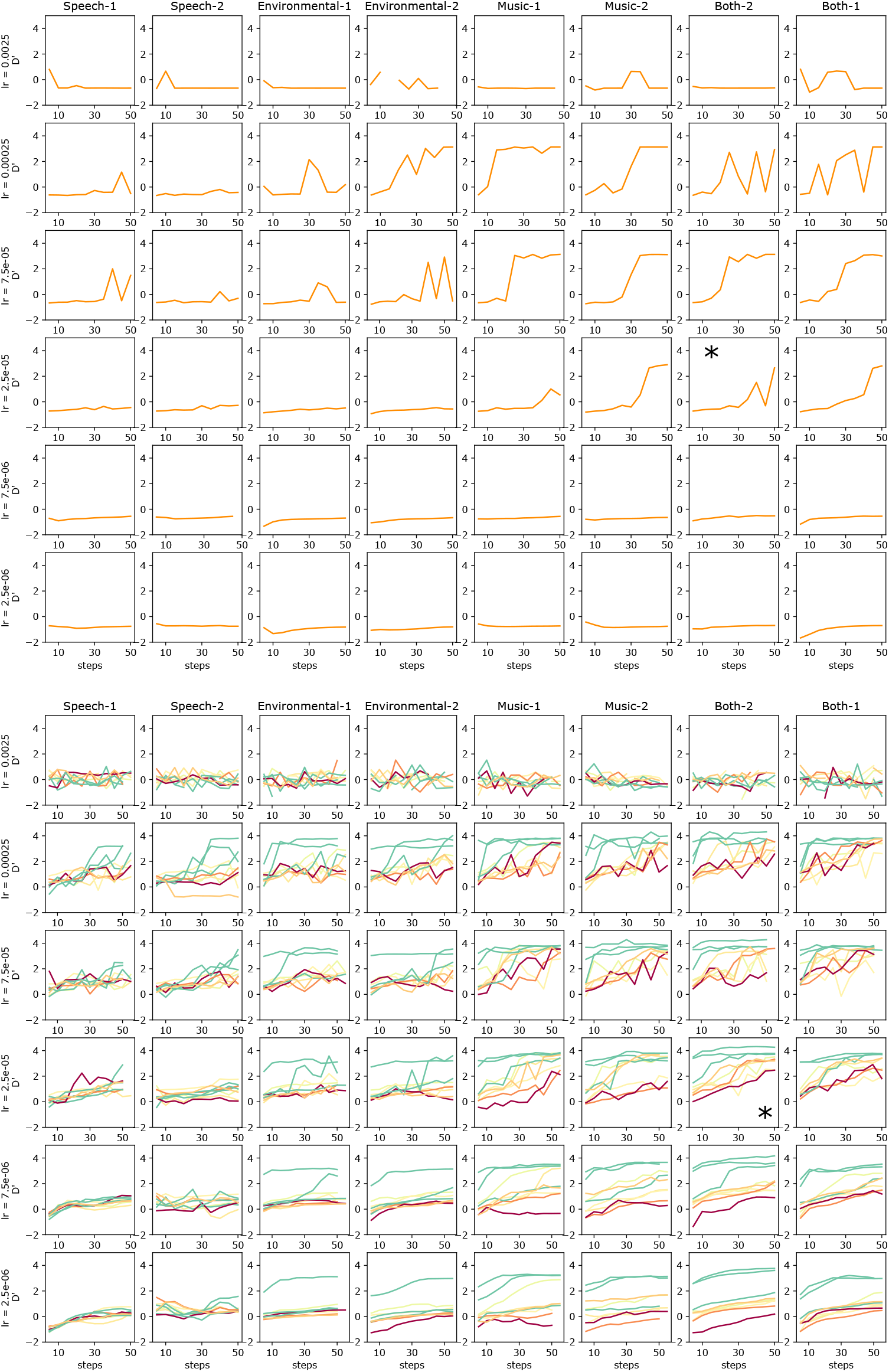
Structure detection in the few-shot scenario, across learning rates (lr) and models. Top: sensitivity to the Global deviant - local standard of the local global paradigm of (18). Note how the performance improve with the learning rate but still requires more than 20 steps of learning. The star indicates the model and learning rate plotted in the supplementary figure 10, with the difference that in supplementary figure 10, the operation was repeated for the last 10 pretraining checkpoints and the d’ averaged across this checkpoint. Here, we only measure and report the d’ for the final checkpoint. Bottom: sensitivity to the deviants in the algebraic pattern paradigms of (16). Note that these evaluations are performed only once per sequence.The fragility and randomness of the model behavior in the few-shot scenario is revealed by the non-monotonous aspect of the curves.

**Supplementary Figure 12.**
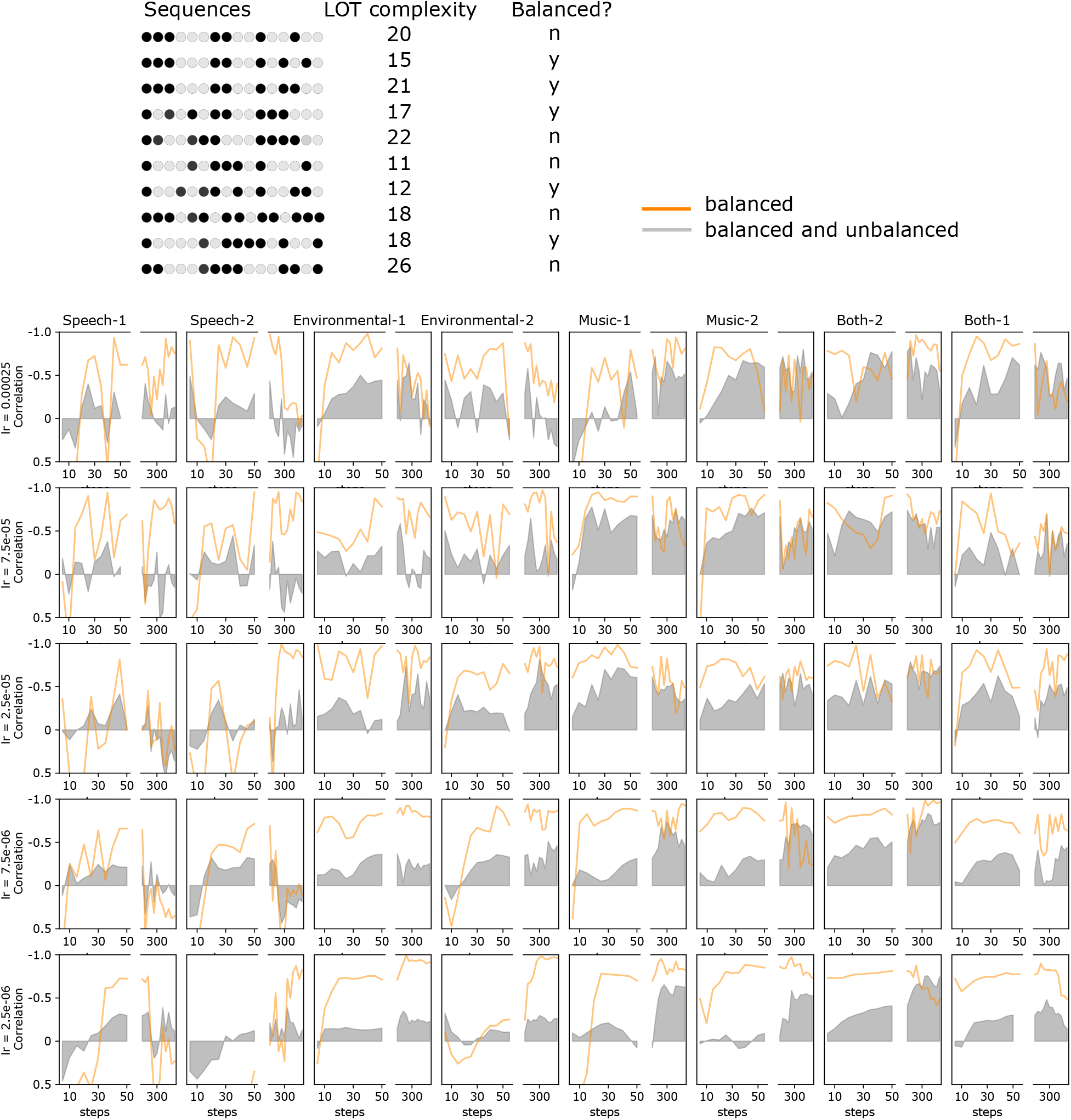
Correlations of the LOT complexity and the model sensitivity for 10 novel sequences of cycle 16 but varied complexity, either balanced as in algebraic patterns of (16), or imbalanced as in the local global paradigm. We plot in grey the correlations of the surprise to all sequences and LOT complexity, and in orange the correlation of the surprise of the 5 balanced sequences and LOT complexity. This correlation is plotted as a function of the habituation steps. The model is modestly correlated with the LOT complexity for most models except speech models where the correlation remains low. This correlation is further improved by using only balanced sequences in most of the case. Once more, we note that in this few-shot scenario, the model abilities are fragile: they vary with the learning rate and are non-monotonously increasing with the steps of habituation.

**Supplementary Figure 13.**
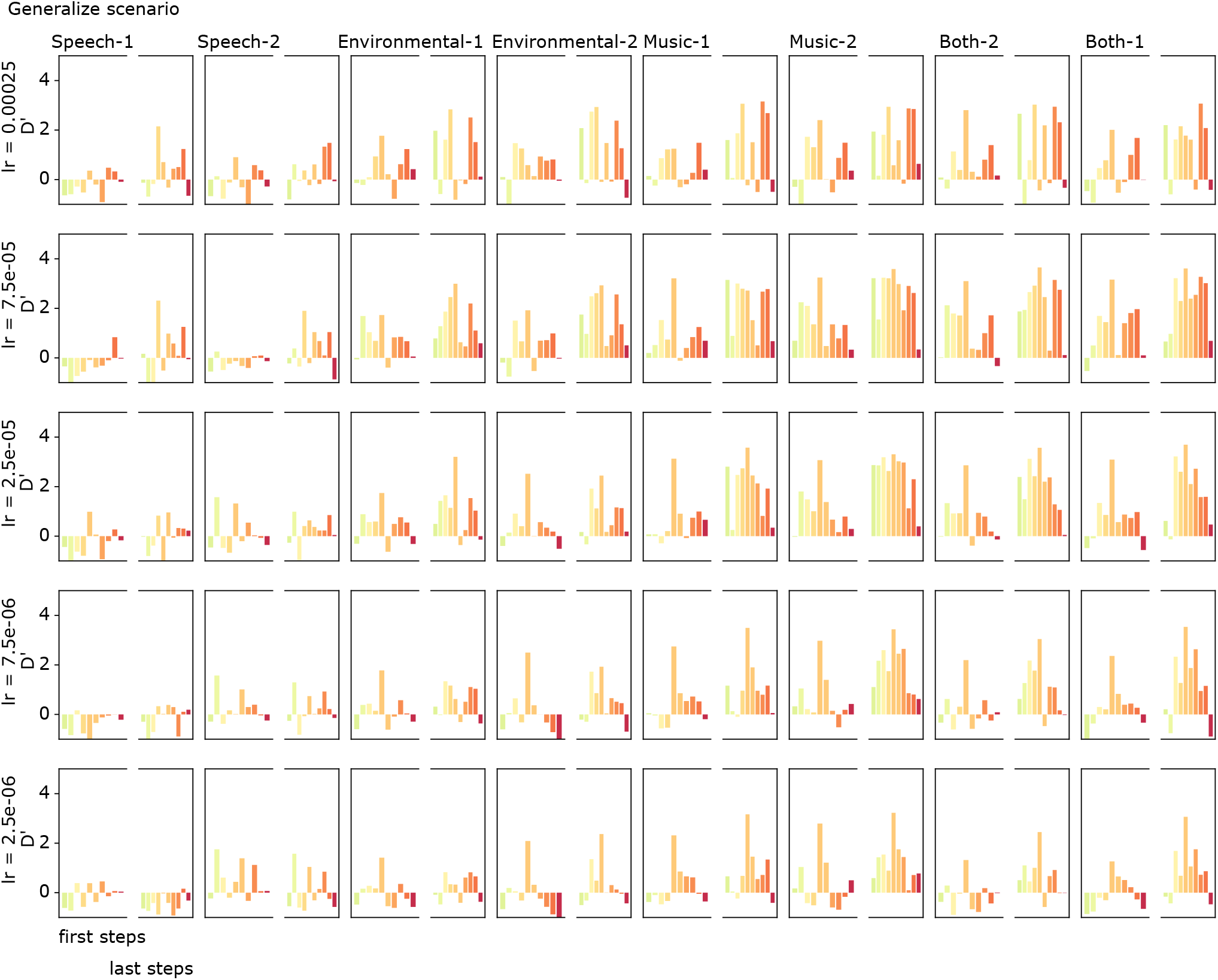
Test of the ability to generalize the algebraic patterns, with ISI = 250 ms. We measure the average model sensitivity across the early 5 to 50 steps of exposition (left), or the late 50 to 600 steps of exposition (rigth), for all models and 5 different learning rates (lr). The models are tested on the novel set of 10 sequences presented in supplementary figure 13, but in the case of the generalization scenario introduced in supplementary figure 5.

## Footnotes

1 The contrastive loss actually compares a projection of the final transformer layer to a discrete encoding of the masked latent. To perform this discrete encoding, code-vectors are learned and the output of the convolutions projected to define a probability of being one of the code-vector

2 As well as additional regularization losses defined in the original Wav2vec2 paper (36)

3 https://github.com/hadware/voxpopuli

## Notes

### Competing Interest Statement

The authors have declared no competing interest.

